# An Atlas of Adaptive Evolution in Endemic Human Viruses

**DOI:** 10.1101/2023.05.19.541367

**Authors:** Kathryn E. Kistler, Trevor Bedford

## Abstract

Through antigenic evolution, viruses like seasonal influenza evade recognition by neutralizing antibodies elicited by previous infection or vaccination. This means that a person with antibodies well-tuned to an initial infection will not be protected against the same virus years later and that vaccine-mediated protection will decay. It is not fully understood which of the many endemic human viruses evolve in this fashion. To expand that knowledge, we assess adaptive evolution across the viral genome in 28 endemic viruses, spanning a wide range of viral families and transmission modes. We find that surface proteins consistently show the highest rates of adaptation, and estimate that ten viruses in this panel undergo antigenic evolution to selectively fix mutations that enable the virus to escape recognition by prior immunity. We compare overall rates of amino acid substitution between these antigenically-evolving viruses and SARS-CoV-2, showing that SARS-CoV-2 viruses are accumulating protein-coding changes at substantially faster rates than these endemic viruses.

## Introduction

Because of their fast mutation rates and high offspring production, many viruses are capable of rapidly evolving to persist and thrive in a changing environment. In the context of human health and disease, this rapid evolution means that viruses from a different host species that cause sporadic human infections can sometimes optimize their cell entry, replication, and immune evasion quickly enough to spread from human-to-human and become a novel pathogen. Thus, the early stages of a pandemic are often marked by high rates of adaptive evolution, as was noted for the 2009 H1N1 pandemic [32, 45, 24], and during the emergence and spread of ‘variant’ viruses during the SARS-CoV-2 pandemic in late 2020 and early 2021 [47].

After this initial adaptation to a new host, some viruses find a niche as an endemic virus where they are able to infect, replicate in, and transmit between humans without continuous adaptive evolution. However, other endemic viruses continue to evolve adaptively [42, 39, 13]. A well-recognized form of this continuing adaptation is antigenic evolution, where the virus and the human adaptive immune system engage in a back-and-forth evolutionary battle — the immune system to neutralize the virus, and the virus to evade neutralization. Viruses that evolve antigenically are particularly capable of causing repeat infections and escaping vaccine-mediated immunity [8]. Therefore, understanding which viruses evolve in this manner is highly relevant for managing viral transmission and mitigating human disease.

Antigenic evolution is a well-noted phenomenon of influenza A/H3N2, where this type of evolution necessitates nearly yearly reformulations of the seasonal influenza vaccine [42, 25, 28]. Serological testing has also demonstrated antigenic evolution in influenza viruses A/H1N1pdm, B/Victoria, and B/Yamagata [3, 18], as well as in seasonal coronavirus 229E [13] and SARS-CoV-2 [50, 1, 12]. On the contrary, measles [46, 34] and influenza C viruses [30] are known to be antigenically stable and do not undergo this mode of continual adaptive evolution. Whether other endemic human pathogenic viruses evolve antigenically is less well understood.

Here, we aim to survey the potential for antigenic evolution across a broad diversity of endemic human viruses. To do this, we use sequencing data to estimate rates of adaptive evolution across each gene in the genome of 28 viruses, which span 10 viral families and a variety of modes of human-to-human transmission. We identify potential antigenically-evolving viruses as those with high rates of adaptation in the protein that mediates receptor-binding, as this is a primary location of antibody neutralization and the locus of antigenic escape mutations in seasonal influenza viruses [52, 49, 25, 9], coronavirus 229E [13] and SARS-CoV-2 [51, 29, 27, 15].

We estimate rates of adaptive evolution from the genetic sequences of viral isolates that have been sampled over time using a McDonald-Kreitman-based method [31, 43] that was formulated for analyzing RNA viruses by Williamson [53], and later, Bhatt et al [7, 6], and then further improved in this manuscript to account for repeated mutations at the same nucleotide position. By estimating rates of adaptation in units of adaptive mutations per codon per year, this method allows us to directly compare adaptive evolution both across the genes of a genome and between different viruses. We find that, in addition to seasonal influenza A and B viruses, norovirus, respiratory syncytial virus (RSV-A and -B), two seasonal coronaviruses (229E and OC43-A), and enterovirus D68 all have elevated rates of adaptation in their receptor-binding proteins, indicating potential antigenic evolution in these viruses. Our results not only increase our understanding of ongoing adaptive evolution in current endemic viruses, but also provide an expectation of antigenic evolution in other related viruses, including future pandemic viruses. In addition to this manuscript, we have made our results viewable at blab.github.io/atlas-of-viral-adaptation/, where interactive plots allow the user to investigate the results as either a comparison of different viruses or as a comparison across the genome of a single virus.

## Results

### An extension of the McDonald-Kreitman method for estimating rates of adaptation in viral genomes

Viruses that undergo antigenic evolution repeatedly evade detection by host antibodies that were elicited by prior infection or vaccination. Under this type of evolution, mutations that alter viral proteins to escape neutralization while retaining necessary viral functions are under positive selection. Thus, antigenic evolution causes the viral genome to continually fix nonsynonymous mutations in epitopes and can be detected as a high rate of adaptive evolution in genes encoding the targets of neutralizing antibodies.

To identify endemic human viruses that are evolving antigenically, we calculated rates of adaptation across the genomes of a wide diversity of viruses using an extension of the McDonald-Kreitman test [31, 53]. This method divides an alignment of viral sequences into temporal windows, and compares the isolates in each window to a fixed outgroup, which represents the historical genome sequence of that virus [7, 6]. The number of adaptive mutations in each time window are calculated as an excess of fixed (or nearly fixed) nonsynonymous mutations above the neutral expectation. The rate of adaptation is then computed as the slope of the linear regression fitting adaptive mutations versus time.

However, because this method uses a fixed outgroup sequence, multiple mutations occurring within the same codon over time can give inaccurate results for a couple reasons. Firstly, whether a mutation is synonymous or nonsynonymous is determined by substituting that mutation into the outgroup sequence. This can result in false assignment if a mutation within that codon has fixed previously. Additionally, repeated mutations at the same nucleotide position will be counted as only a single mutation because the method has no ‘knowledge’ of previous time windows. This flaw will cause a disproportionate underestimation of the rate of adaptation in viral proteins where many sites have sequentially fixed multiple mutations, as occurs in rapidly-evolving viruses such as influenza A/H3N2 (Figure 1A).

**Figure 1:**
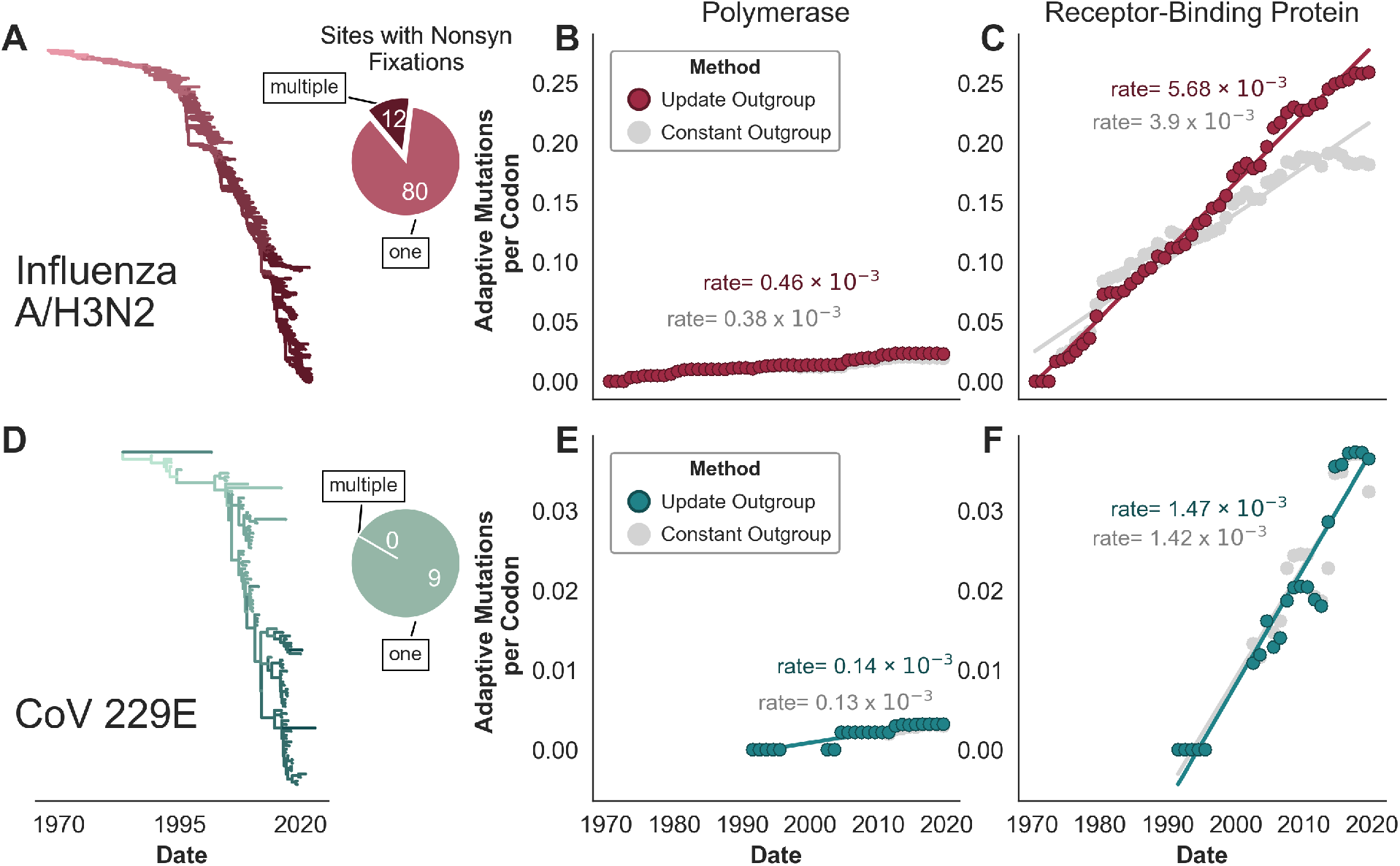
A McDonald-Kreitman-based method to estimate the rate of adaptation in antigenically-evolving viruses. **A)** Time-resolved phylogeny of 2104 influenza A/H3N2 HA sequences sampled between 1968 and 2022 and colored by nonsynonymous mutation accumulation from the root, with darker reds symbolizing more mutations in the HA1 subunit. Within these samples, ninety-two nucleotide sites have completely fixed a nonsynonymous mutation, and the pie chart indicates that 12 of these nucleotide sites have fixed multiple nonsynonymous mutations during the past *∼*50 years. **B)** Accumulation of adaptive mutations (per codon) in polymerase PB1 as calculated by the McDonald-Kreitman-based method that updates the outgroup sequence at each fixation (dark red), or uses a constant outgroup sequence (gray). The rate of adaptation is the slope of the linear regression fitting these estimates. **C)** Estimated accumulation of adaptive mutations in HA1. **D)** Time-resolved phylogeny of 95 coronavirus 229E Spike S1 sequences sampled between 1989 and 2022, colored as in panel A. Pie chart indicates that, within these samples, nine nucleotide sites have completely fixed a nonsynonymous mutation and zero nucleotide sites have fixed multiple nonsynonymous mutations. Accumulation of adaptive mutations, as in panels B and C, within the coronavirus 229E **E)** polymerase (RdRp) and **F)** receptor-binding subunit S1.

To address both of these issues, we have modified the method to update the outgroup sequence each time a mutation fixes. Isolates in later time windows are, thus, ‘aware’ of any fixations that occurred in the same codon during previous time windows. This modification substantially affects the rate of adaptation in the influenza A/H3N2 HA1 subunit, where 92 nucleotide sites have fixed nonsynonymous mutations since 1968 and 12 of these sites have seen multiple nonsynonymous fixations (Figure 1A). The asymptoting shape of the inferred accumulation of adaptive mutations in H3N2 HA1 reflects saturation where many adaptive mutations at later time windows occur at the same position as adaptive mutations that occurred in previous time windows (Figure 1C). In contrast to H3N2 HA1, there are zero nucleotide sites within the S1 subunit of seasonal coronavirus 229E that have fixed multiple nonsynonymous mutations. Fittingly, updating the outgroup sequence has little-to-no effect on the estimated rates of adaptation in the 229E receptor-binding subunit S1 (Figure 1F).

Influenza A/H3N2 [49, 25] and coronavirus 229E [13] are both known to undergo antigenic evolution through the fixation of mutations in their receptor-binding proteins. Because one goal of this manuscript is to quantitatively compare antigenic evolution between viruses by estimating rates of adaptive evolution in the receptor-binding proteins, we use the method that updates the outgroup sequence throughout this study as it better captures the rate of a rapidly-adapting protein, such as H3N2 HA1. However, it should be noted that the major findings and themes presented here do not depend on which version of the method is used — both methods identify the same subset of viruses as antigenically evolving, though the relative pace of this evolution is dependent on which method is used.

### Estimation of the threshold of antigenic evolution

Antigenic evolution occurs when a virus fixes mutations at or near sites of antibody binding that abrogate those antibodies’ abilities to neutralize the virus. For viruses that have been demonstrated to evolve antigenically, these escape mutations occur in the viral protein that mediates receptor-binding, which is located on the virion’s surface and is typically a primary target of neutralizing antibody binding. For instance, in influenza A/H3N2, antigenic evolution occurs largely in HA1, the receptor-binding subunit of hemagglutinin [52, 49, 25, 9]. Similarly, for seasonal coronavirus 229E, escape mutations fix in S1, the receptor-binding subunit of Spike [13]. Thus, we hypothesize that antigenic evolution results in a high rate of adaptation in the receptor-binding protein or subunit, and that antigenically-evolving viruses can be distinguished from antigenically-stable ones by the rate of adaptation on the receptor-binding protein.

We calculated the rate of adaptive evolution in the receptor-binding protein for three viruses which are known to evolve antigenically — influenza viruses A/H3N2 and B/Yam [3] and coronavirus 229E [13] — as well as for three viruses that are known to be antigenically stable — measles [46, 34], influenza C/Yamagata [30], and hepatitis A [16, 2]. All of the antigenically-evolving viruses have higher rates of adaptation than the antigenically stable viruses (Figure 2A), indicating that this method successfully differentiates between viruses that evolve antigenically and those that do not. We used these rates of adaptive evolution to estimate a threshold of antigenic evolution (i.e. a rate above which we predict the virus to be evolving antigenically) using logistic regression (see Methods for more details). We estimated that the threshold of antigenic evolution is about 1.18 × 10^−3^ mutations per codon per year in the receptor-binding protein. We view this threshold not as an absolute, but rather, as our best approximation of a rate that separates evolution with real biological meaning from noise in the estimates.

**Figure 2:**
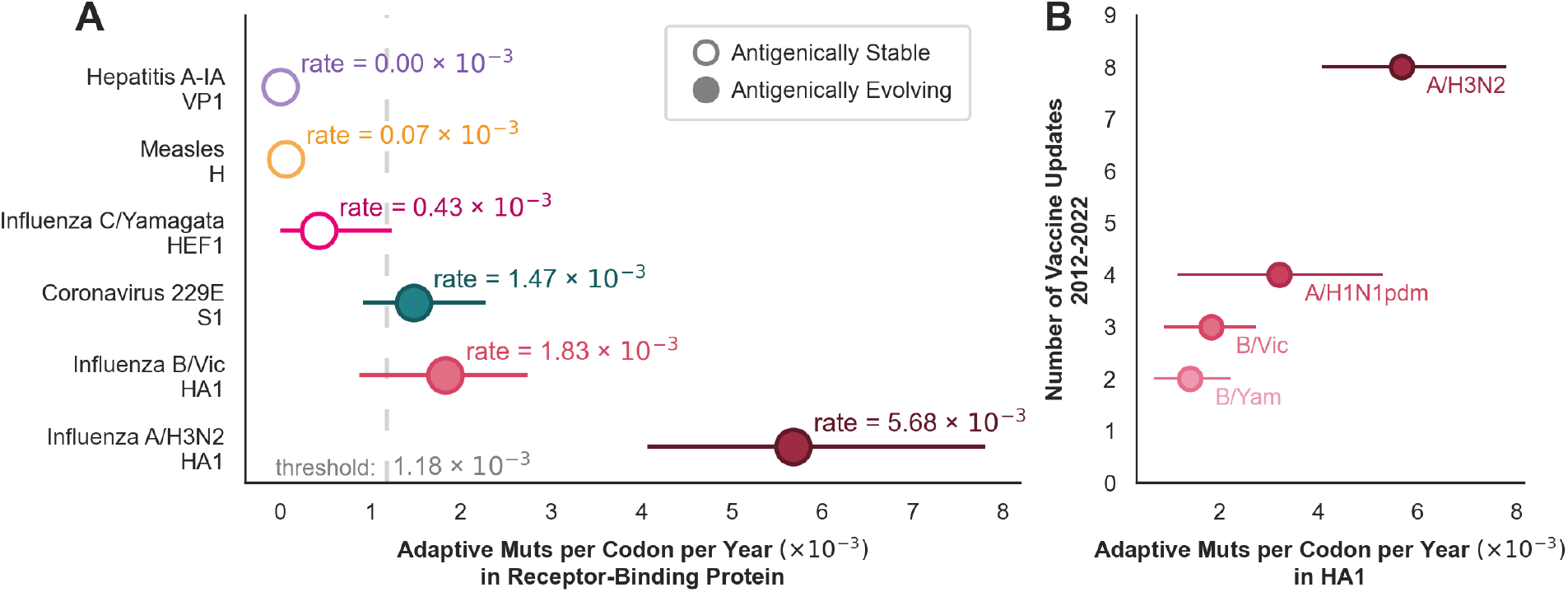
Rates of adaptation in the receptor-binding protein recapitulate known trends of antigenic evolution. **A)** The rate of adaptation calculated in the receptor-binding protein is plotted for 3 antigenically-stable viruses (solid circles), and 3 antigenically-evolving viruses (open circles). The threshold of antigenic evolution is estimated by logistic regression. Error bars represent the 95% bootstrap percentiles. **B)** For each of the 4 influenza viruses that are included in the yearly flu vaccine, the rate of adaptation is compared with the number of times that the vaccine strain was updated between the 2012-2013 and 2022-2023 Northern hemisphere flu seasons.

The relative rates of adaptation estimated here are consistent with what is already known about the relative pace of antigenic evolution in these viruses. Bedford et al [3] estimated that influenza A/H3N2 evolves antigenically 2-3 times faster than the influenza B viruses, and Eguia et al [13] found that coronavirus 229E escapes neutralization at a rate similar to influenza B. Additionally, our estimated rates of adaptation in HA1 of the various influenza viruses reflect the frequencies at which these viruses deviate antigenically enough to warrant an update to the vaccine strain (Figure 2B). Influenza A/H3N2 exhibits the highest rate of adaptation (5.7 × 10^−3^ mutations per codon per year), followed by A/H1N1pdm (3.2 × 10^−3^ mutations per codon per year), then B/Vic (1.8 × 10^−3^ mutations per codon per year) and B/Yam (1.4 × 10^−3^ mutations per codon per year). Mirroring this, the A/H3N2 component of the vaccine has been updated 8 times (9 different strains) between 2012 and 2022, while the H1N1 strain was updated 4 times, the B/Vic component was updated 3 times, and B/Yam component was updated twice during this time period [54]. Components of the seasonal influenza vaccine are updated by the World Health Organization (WHO) Global Influenza Surveillance and Response System (GISRS) when the vaccine strain no longer induces sufficient protection against circulating viruses — a point which is typically defined by an 8-fold drop in titer in a hemagglutinin inhibition (HI) assay [55]. This suggests that the rate of adaptation in the receptor-binding domain can identify not just which viruses evolve antigenically, but also the relative pace at which they do so and, thus, the expected duration of protection afforded by antibodies elicited by vaccination or infection.

### Genome-wide appraisal of rapidly-evolving viral proteins

We next sought to survey a wide-diversity of endemic human viruses for evidence of ongoing adaptive evolution during the past *∼*50 years. To do this, we downloaded and curated sequence data for 28 human pathogenic viruses, which belong to 10 different viral families. This panel comprises both RNA and DNA viruses with a variety of modes of transmission including respiratory, fecal-oral, vector-borne, and via blood or bodily fluids. We focus on viruses that have been endemic in humans for at least 12 years because we are interested in continued adaptive evolution that persists during the endemic phase (rather than initial host adaptation that occurs early in a pandemic) and because we have previously shown that a short temporal spread of sampled sequences decreases the accuracy of the estimated rate of adaptation [23].

For each virus, we estimated the rate of adaptation in each gene of the genome. In total, we analyzed 239 viral genes and 14 of them had rates of adaptation exceeding the threshold of antigenic evolution (Figure 3A). Of these 14 genes, 13 encode proteins located on the viral surface, with 10 of those being proteins that mediate host receptor-binding (Figure 3B). The 3 surface proteins with rates of adaptation exceeding our threshold that are not classified as receptor-binding are all neuraminadase (NA) genes of the influenza subtypes A/H3N2, A/H1N1pdm, and B/Vic. The influenza NA protein has been shown to bind host receptors in some influenza viruses [56, 21], and to be a target of protective and neutralizing antibodies [10, 14, 44], so it is possible that these high rates of adaptation in influenza NA are also reflective of evolution to escape antibody recognition. The non-surface protein that has a high rate of adaptation is norovirus p22, which antagonizes cellular protein trafficking [41], and has been previously reported to be under positive selection [11].

**Figure 3:**
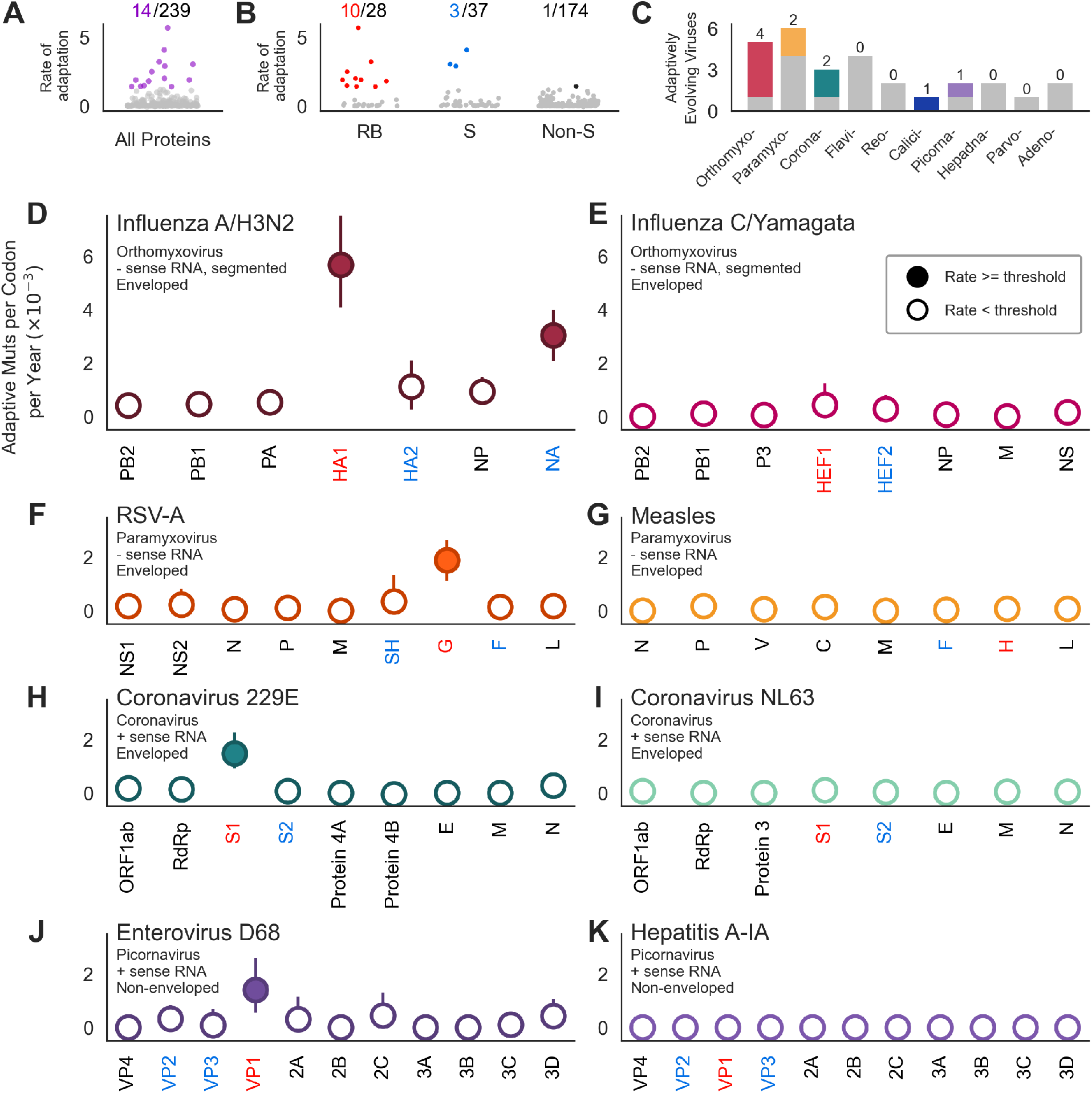
Across 28 viral genomes, the highest rates of adaptation are found in surface-located receptor-binding proteins. **A)** The rate of adaptation for all 239 viral genes. Fourteen genes (in purple) have rates of adaptation above our threshold of antigenic evolution. Genes with rates of adaptation below the threshold are in gray. **B)** The rate of adaptation within all 28 receptor-binding proteins (RB, left), 37 other proteins located on the viral surface (S, center), and 174 non-surface proteins (Non-S, right). Ten receptor-binding proteins (red), 3 other surface-located proteins (blue) and 1 non-surface protein (black) exceed our threshold. Genes with rates below the threshold are in gray. **C)** Number of viruses per viral family that have at least one gene exceeding the threshold are shown in color. The number of viruses in these families that had no high rates of adaptation throughout their entire genome is in gray. **D-K)** Rates of adaptation were calculated for each gene, subunit, or coding region indicated along the x-axis, and ordered by genomic position (or segment number, for segmented viruses). Receptor-binding proteins are labeled in red, other surface-exposed proteins are in blue, and non-surface-located proteins are in black. Filled circles indicate genes with rates exceeding the threshold. Each row shows two viruses from the same viral family, one which contains at least one adaptively-evolving gene (left) and one which does not (right). Error bars indicate the 95% bootstrap percentiles from 100 bootstrapped data sets.

In total, 10 of the 28 viruses in this panel had at least one gene we predict to be undergoing ongoing adaptation. These viruses include members of the orthomyxovirus (influenza A/H3N2, A/H1N1pdm, B/Vic and B/Yam), paramyxovirus (RSV-A and RSV-B), coronavirus (229E and OC43-A), calicivirus (norovirus GII.4), and picornavirus (enterovirus D-68) families (Figure 3C). While multiple orthomyxo-, corona- and paramyxoviruses appear in this list, our results suggest adaptive evolution is not necessarily a shared feature of related viruses. For instance, while influenza A/H3N2 has two adaptively evolving proteins, influenza C/Yamagata has none (Figure 3D and E). Similarly, 229E and NL63 are both alphacoronaviruses, but 229E has a high rate of adaptation in spike S1, while NL63 does not (Figure 3H and I).

Among the 10 viruses with at least one protein exceeding our threshold, we observe that highest rates of adaptation genome-wide are typically the genes encoding the receptor-binding protein or subunit. The exceptions being influenza A/H1N1pdm and B/Vic, where the fastest rates are in NA not HA1-though, as mentioned above, NA is sometimes involved in receptor-binding. These results reveal that endemic viruses experience little-to-no ongoing adaptation throughout most of their genome and that continuous adaptive evolution is found almost solely in surface-exposed proteins, which are accessible to neutralizing antibodies. This suggests that evasion of antibody neutralization is a driving force in the ongoing adaptive evolution of many endemic viruses.

### Identification of putative antigenically-evolving viruses

With the expectation that antigenic evolution is detectable by a high rate of adaptation in the receptor-binding protein, we then directly compared rates between the receptor-binding proteins of 28 viruses (Figure 4). We also compared the rates of adaptation in the polymerase, which, in endemic viruses, we expect to be relatively conserved. We observe that while there is little variation between the rates of adaptation in the polymerase, which range between 0.0 and 0.7 × 10^−3^ mutations per codon per year, there is much larger spread of rates in the receptor-binding proteins of these viruses (ranging from 0.0 to 5.7 × 10^−3^ mutations per codon per year). Based on the estimated rates of adaptation that exceed the threshold for antigenic evolution we identified, we predict that 10 of the viruses in this panel evolve antigenically.

**Figure 4:**
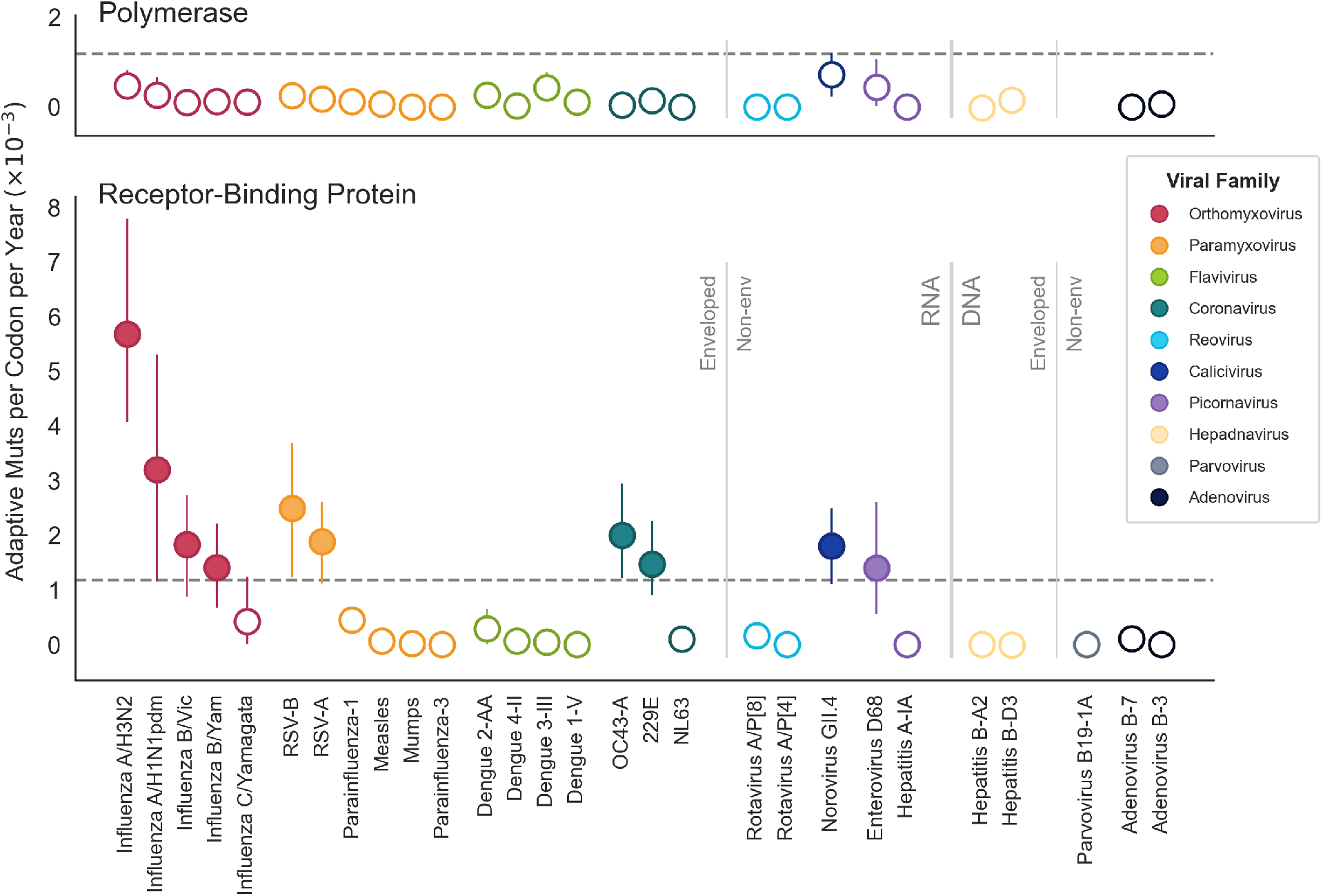
Comparison of rates of predicted antigenic evolution across a wide-diversity of human pathogenic viruses. Rates of adaptive evolution in the polymerase (top) and receptor-binding protein (bottom) for 28 human pathogenic viruses. The threshold of antigenic evolution (as determined in Figure 2) is marked by the dotted line; the rates falling above this line are shown by solid markers, and the rates below the threshold are open circles. Viruses are grouped and colored by viral family, and arranged within viral family in descending order of the receptor-binding rate. Viral families are ordered by genome type, with RNA viruses shown in brighter colors and DNA viruses in graytones. Vertical dividers further delineate enveloped from non-enveloped viruses.

Influenza A/H3N2 has, by far, the fastest rate of antigenic evolution, followed by A/H1N1pdm, while the other 8 putative antigenically-evolving viruses all have rates in roughly the same range, around 1.5 − 2 × 10^−3^ adaptive mutations per codon per year. In descending order of the estimated rate of antigenic evolution, these viruses are: RSV-B, RSV-A, norovirus GII.4, coronavirus OC43-A, enterovirus D-68, influenza B/Yam, coronavirus 229E, and influenza B/Vic.

All of these 10 viruses have RNA genomes, though both positive- and negative-sense genomes and both enveloped and non-enveloped viruses appear capable of antigenic evolution. All of these viruses transmit via a respiratory route except norovirus, which uses fecal-oral transmission. We observe that multiple coronaviruses, RSV viruses, and influenza viruses evolve antigenically, suggesting these types of viruses might have a higher propensity for this type of evolution. However, in each of these cases, we also find that at least one other member of the same viral family does not evolve antigenically, indicating that relatedness at the level of viral family is not an absolute predictor of antigenic evolution. Overall, the examination of antigenic evolution presented here suggests that selection to evade antibody recognition is widespread, though certainly not ubiquitous, among endemic RNA viruses, and that, while certain types of viruses may have a higher propensity for this type of evolution, even closely-related viruses can differ in this regard.

A comparison of rates of adaptation in the receptor-binding proteins of all the viruses in this panel, as well as rates across the genome of each of these viruses can be viewed interactively at blab.github.io/atlas-of-viral-adaptation/ (see example screenshots in Supplementary Figure S1). This interactive website shows the rates per codon per year (as reported in this manuscript), as well as per gene per year, allows the viruses to be ordered by rate rather than by viral family, and displays rates calculated both by the constant outgroup and updated outgroup methods.

### Comparison of rates of evolution between endemic viruses and SARS-CoV-2

An obvious question is where the evolution of SARS-CoV-2 falls with respect to these other viruses. Since the beginning of the SARS-CoV-2 pandemic, we have seen a period of many co-circulating variants that contained adaptive mutations, followed by a single fixation event where Omicron swept, and then a subsequent period where many competing Omicron lineages are co-circulating, with repeated near sweeps of derived lineages such as BA.5, BQ.1 and XBB.1.5 that are supplanted before reaching fixation. Because the McDonald-Kreitman-based rate estimation we have used thus far considers only fixed or nearly-fixed mutations to be potentially adaptive, this method is ill-suited to analyzing SARS-CoV-2 evolution so far. Essentially, the calculated rate will just reflect the high number of mutations on the long branch leading to Omicron that fixed when Omicron swept. Additionally, this method can be noisy over short time periods, where small numbers of fixations can have an outsized effect on the rate. It is for this reason that we limited our panel to viruses that have been endemic for several years, with influenza A/H1N1pdm having the narrowest span of human circulation (12 years).

In lieu of calculating a rate of adaptation for SARS-CoV-2, we instead do a much simpler comparison of the rates of amino acid substitution in the receptor-binding proteins between SARS-CoV-2 and the 10 viruses we predict to be evolving antigenically (Figure 5). We find that SARS-CoV-2 accumulates roughly 20 × 10^−3^ amino acid substitutions per residue per year in S1. This is 2-2.5X faster than the accumulation of amino acid substitutions in influenza A/H3N2 HA1 and 7-10X faster than in the S1 subunit of seasonal coronaviruses 229E and OC43. Importantly, the rate of amino acid substitution among all SARS-CoV-2 viruses is not solely driven by the fixation of 12 S1 substitutions when Omicron swept; in fact we observe roughly the same as the rate between SARS-CoV-2 viruses spanning the entire pandemic (Figure 5A) as we do among the currently predominant clade of Omicron 21L and its descendants (Figure 5B), corresponding to lineage BA.2 and derived lineages like BQ.1 and XBB.

**Figure 5:**
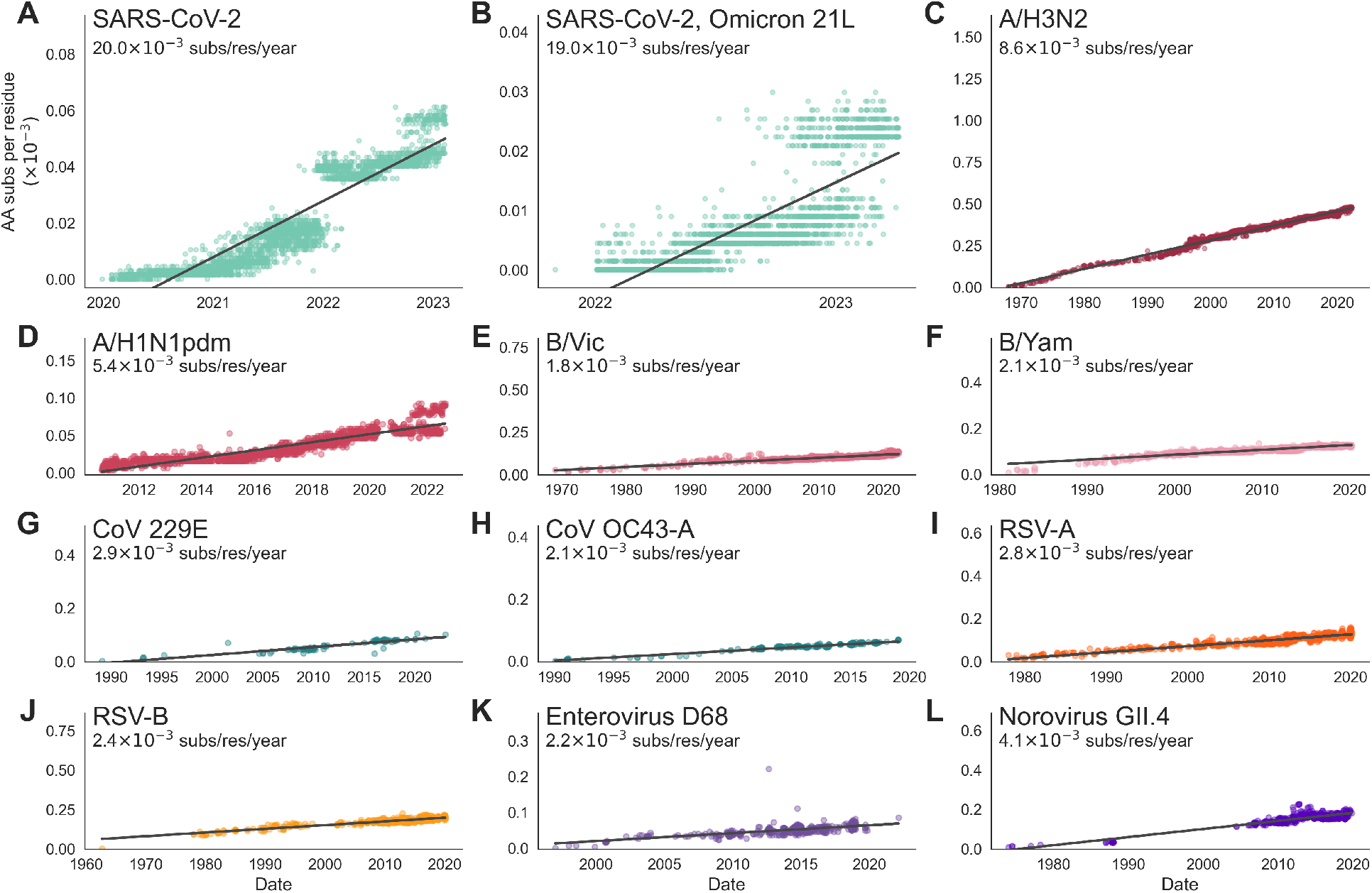
Rates of amino acid substitution in the receptor-binding protein of SARS-CoV-2 and 10 antigenically-evolving endemic viruses. The rate of amino acid substitution in the receptor-binding protein of **A)** SARS-CoV-2, **B)** SARS-CoV-2 Omicron clade 21L, **C)** influenza A/H3N2, **D)** influenza A/H1N1pdm, **E)** influenza B/Vic, **F)** influenza B/Yam, **G)** coronavirus 229E, **H)** coronavirus OC43-A, **I)** RSV-A, **J)** RSV-B, **K)** enterovirus D68, **L)** norovirus GII.4. Rates are computed as the slope of a linear regression fitting a comparison of amino acid substitutions versus time, and are found using a phylogeny. Each tip on the tree is plotted by its sampling date and the number of amino acid substitutions that accumulated between the root and the tip (normalized by the length of the coding region, in residues). Aspect ratios in each panel are fixed so that regression slopes are visually comparable across panels.

In Figure 5, we simply plot the number of amino acid changes in the receptor-binding protein that each tip has compared to the root. This simpler analysis does not consider fixation of particular mutations, nor does it make any attempt to account for substitutions under selection versus those that are present due to chance or hitch-hiking. However, we find that this simpler analysis reflects the general relationships between rates of antigenic evolution of different viruses that we present in Figure 4. Figure S2 lists the rate of amino acid substitution and the rate of adaptation in the receptor-binding protein of each virus. A ratio of the rates in Figure 4 and Figure 5 indicates that, in most antigenically-evolving endemic viruses, between *∼*60 and 100% of amino acid substitutions in the receptor-binding protein are adaptive. For instance, we estimate that influenza A/H3N2 evolves antigenically at 5.7 × 10^−3^ adaptive mutations per codon per year, which is *∼*66% of the 8.6 × 10^−3^ amino acid substitutions per residue it accumulates each year. Of the 10 antigenically evolving endemic viruses, the lowest proportion of amino acid substitutions that are adaptive is 44% in norovirus GII.4 VP1. Thus, if we presume that a similar number of SARS-CoV-2 amino acid substitutions are adaptive, we get a rate of between *∼*9 and *∼*20 × 10^−3^ adaptive mutations per year in S1.

## Discussion

In search of antigenically-evolving viruses, we use genomic sequences to analyze adaptive evolution across a panel of 28 viruses and present the results here and as a website with interactive plots and phylogenies at blab.github.io/atlas-of-viral-adaptation/. We find that antigenic evolution is not uncommon amongst endemic human viruses, with ten viruses spanning five viral families meeting our criteria for predicting antigenic evolution. Particularly, this mode of evolution seems prevalent amongst endemic viruses that have RNA genomes (Figure 4). However, the true proportion of endemic viruses that evolve antigenically is hard to estimate because the panel of viruses analyzed here is far from a comprehensive list of human endemic viruses and is biased towards well-studied viruses, which are not necessarily the most common pathogens. We selected viruses to include in the panel based on the following criteria: 1) virus has been endemic for at least 12 years, 2) the genome is under 50 kilobases, and 3) there are at least 50 high-quality genomes available spanning at least 12 years. For many endemic viruses, the limiting factor is a dearth of historical sequences predating the mid-2000’s. However, the COVID-19 pandemic has spurred an increased interest in monitoring and sequencing human pathogens and, if this trend continues, it is likely that there will be enough longitudinal data to add many more viruses to this panel in the years to come.

By employing a quantitative method, we are able to compare the pace of adaptive evolution between genes in a genome as well as between viruses. Comparisons within genomes reveal that surface proteins are consistently the fasting-evolving viral proteins (Figure 3). Comparisons between viruses show that influenza A/H3N2 is especially striking its pace of antigenic evolution, which is roughly 2-3 times faster than all other viruses we predict to be evolving antigenically (Figure 4). The observation that many viruses accumulate adaptive mutations in their receptor-binding protein at a rate of roughly 1.5 − 2.0 × 10^−3^ mutations per codon per year suggests that this might be a lower bound on the rate that is sufficient to generate antigenic novelty fast enough that a virus can persist in a mostly-immune population. However, it should be stressed that while the rate of adaptation is similar between these eight viruses, it is not clear whether the pace of relevant phenotypic change is also similar between them. For instance, it may be that some viruses need, say, two adaptive mutations on average to successfully escape prior immunity, while other viruses only need one.

Whether or not a virus evolves antigenically, and the pace at which is does so, is likely a function of many factors, including mutation rate, mutational tolerance of surface proteins [48], positioning and co-dominance of epitopes [34], viral transmission dynamics, and existing population immunity, to name a few. Our estimates of rates of antigenic evolution do not allow us to disentangle which factors are contributing most to the evolution of endemic viral proteins. However, it is an interesting and open question why closely-related viruses (such as coronaviruses 229E and NL63) differ in their propensity to evolve evasion of antibody detection and, relatedly, what the minimal necessary information to predict this type of evolution in an emerging virus is.

In this study, we have focused on continuous antigenic evolution within viral lineages over the past *∼*50 years. It is important to note that this is a very particular type of evolution in which antigenic variation is selected for repeatedly, leading to selective sweeps within a single lineage. However, this does not necessarily mean that viruses that are antigenically-stable on a *∼*50-year time scale have not undergone some form of antigenic evolution in the past or at other time-scale. For instance, influenza C viruses and dengue viruses both exist as several antigenically-distinct lineages, and while ongoing antigenic evolution is not occurring *within* these lineages, the establishment of antigenically-distinct lineages was likely a result of selection in the past. It is possible that some viruses are more prone to fracturing into several antigenically-distinct, co-circulating lineages rather than undergoing perpetual antigenic evolution within a lineage.

Relatedly, it is important to note that this method looks for fixations and near fixations, with the idea that positively-selected mutations will sweep through the population. This means that mutations that fix within a clade, but not the entire population, will not be considered potentially-adaptive and, thus, that this method is sensitive to how lineages are designated. For instance, if all influenza B viruses were analyzed together, rather than as separate B/Vic and B/Yam lineages, there would be no signal of adaptation. In some cases it can difficult to define what constitutes two distinct lineages versus two clades of the same lineage. In our analyses, we have divided each viral species into the lineage or genotype classifications used by the field of literature for that virus.

### Implications for the ongoing evolution of SARS-CoV-2

In SARS-CoV-2 S1, we observe a rate of amino acid substitution that is roughly 2-2.5X the rate in influenza A/H3N2 HA1, the prototypical example of rapid antigenic evolution. There is an open question of whether SARS-CoV-2 can sustain such high rates of evolution in the years to come. To address this question, we can retroactively observe how the evolution of other viruses has changed between the early pandemic and the ensuing endemic years. In this manuscript, we have analyzed the evolution of influenza viruses A/H3N2 and A/H1N1pdm over time between their respective introductions in 1968 and 2009, and today. We do not see any evidence that the rate of amino acid substitution (Figure 5C and D) or the rate of adaptive evolution (Figure 1C) in HA1 is flagging in these antigenically-evolving influenza viruses. By extension, this suggests that we may continue to see rapid evolution in the S1 subunit of SARS-CoV-2. This fits with our observation that the rate of amino acid substitution in S1 of Omicron clade 21L viruses circulating in 2022 and 2023 is roughly the same as this rate over entire pandemic (Figure 5A and B), suggesting that, so far, S1 evolution has not slowed throughout the course of the pandemic.

While overall we expect that SARS-CoV-2 will continue to evolve appreciably faster rates than seasonal influenza or coronaviruses, it is unclear whether this evolution will be somewhat slowed by the build-up of increasingly complex immune histories toward this virus. At this point, it is also difficult to predict whether the emergence of a highly fit and highly divergent variant (Omicron) was a one-time event, or whether other similar lineages will emerge in the future and continue to be a feature of SARS-CoV-2 evolution.

That SARS-CoV-2 is able to evolve antigenically has become readily apparent over the three years since the beginning of the pandemic [12, 1, 50, 15]. However, in early 2020, at the beginning of the COVID-19 pandemic, it was not known whether related coronaviruses evolve antigenically and, thus, it was difficult to speculate whether SARS-CoV-2 would evolve in this way or not. We believe this issue reveals how little is known about the antigenic evolution of many of the viruses that commonly infect us. We believe a better understanding of the broad diversity of endemic viruses will not only better prepare us for future pandemics, but also will inform our current efforts to design vaccines and therapeutics against these viruses. To this end, we have compiled this atlas of viral adaptive evolution to quantitatively compare evolution across a wide range of endemic viruses.

## Methods

The code to implement the McDonald-Kreitman-based calculations of adaptation rates is located at https://github.com/blab/adaptive-evolution. All analysis code is written in Python 3 (Python Programming Language, SCR 008394) in Jupyter notebooks (Jupyter-console, RRID:SRC 018414). The results presented in this manuscript are also accessible in an interactive format at https://blab.github.io/atlas-of-viral-adaptation/.

### Input data for the estimation of rate of adaptation

The analyses in this manuscript require an alignment file in FASTA format, a tabular metadata file that contains the sampling date for each sequence in the alignment, and a reference file in Genbank format that supplies gene locations. We have written the analysis code with the intention that it should be paired with a Nextstrain build [17], which will create the necessary alignment and metadata files as well as a phylogenetic tree, which is not necessary for the analysis but is a useful companion for the interpretation of the results. Of the pathogens considered in this manuscript, we used builds produced and maintained by the Nextstrain team for influenza A and B (https://github.com/nextstrain/seasonal-flu, created as part of [35]), measles (https://github.com/nextstrain/measles), mumps (https://github.com/nextstrain/mumps, created as part of [33]), dengue (https://github.com/nextstrain/dengue, created as part of [4]), and enterovirus D68 (https://github.com/nextstrain/enterovirus_d68, created as part of [20]). For the influenza A and B viruses, we tweaked the builds to contain sequences from any date, rather than limiting the tree to isolates sampled within the past 12 years. For dengue, we adjusted the pipeline to produce genotype-level phylogenies, rather than serotype-level. For all other viruses, we constructed a new Nextstrain build, as described below.

### Sequence data

Norovirus, RSV, and rotavirus sequences were downloaded from ViPR/BV-BRC [38, 37]. Adenovirus, hepatitis A, hepatitis B, parvovirus B19 genomes were downloaded from Genbank [5]. Parainfluenza and seasonal coronavirus sequences were downloaded from both Genbank ViPR/BV-BRC and combined. Influenza C sequences were downloaded from NCBI Viruses [19]. All sequence queries were limited to clinical isolates from human hosts. All sequence data for a pathogen was curated into a single FASTA file, excluding sequences that had no available date information. Compiled sequence data for all these viruses are available via https://github.com/blab/adaptive-evolution.

### Nextstrain builds to generate alignments and trees

Time-resolved phylogenies were generated for each pathogen by running a Nextstrain build [17]. Sequences were aligned to a reference genome using MAFFT [22]. Trees were constructed using IQ-TREE [36], and branch lengths were inferred with TreeTime [40]. Builds were streamlined into a pipeline using Snakemake [26], and Snakemake workflows are available for each virus via https://github.com/blab/adaptive-evolution.

### SARS-CoV-2 Nextstrain phylogenies

SARS-CoV-2 alignments and trees were retrieved from nextstrain.org builds. The build spanning all SARS-CoV-2 lineages contains samples from the beginning of the pandemic until February 13, 2023 and contains sequences evenly sampled over time and geography. This dataset is viewable at https://nextstrain.org/ncov/gisaid/global/all-time/ 2023-02-26. The 21L-only build contains only sequences from the Omicron clade 21L up until April 9, 2023. This dataset is viewable at https://nextstrain.org/ncov/gisaid/21L/global/all-time/2023-04-18.

### Rate of adaptation, with a fixed outgroup

The rate of adaptation within each gene of a genome is calculated from alignment of viral sequences sampled over time as in Bhatt et al, 2011 [6]. To do this, the sequence alignment is broken up into constituent genes or subunits. Then, the gene-specific alignment is partitioned into 5-year windows tiling the entire span of time over which data is available. Windows are offset by 1 year so, for example, an alignment containing sequences from 1990-2022 would be partitioned into windows of [1990-1995, 1991-1996, …, 2016-2021, 2017-2022]. The exceptions are H1N1pdm and mumps where we use 3-year windows rather than 5-year, because there are only 12 and 17 years of data, respectively. The window size is a trade-off between picking up more signal (shorter windows), and reducing noise (longer windows) that can be due single sequences having a outsized effects on small sample sizes or chance sampling of one co-circulating clade over another. We require that each temporal window contain at least 3 isolates, and exclude time windows with 2 or fewer samples.

The outgroup sequence is found by taking a consensus of the sequences present in the first window. The choice to use a consensus sequence, rather than Most Recent Common Ancestor (MRCA), as the outgroup was based on previous implementations of this method [53, 6], to keep the method alignment- rather than phylogeny-based, and because, in our intial testing, similar rate estimates were obtained using MRCA or consensus outgroup. Each subsequent temporal window is then compared to the outgroup sequence to find polymorphisms and fixations. To do this, each nucleotide position in the gene alignment is compared to the outgroup to determine polymorphism, fixation, replacement and silent scores (see [7] and [6] for more details). The expectation for neutral evolution is found from the number of polymorphisms present at 15-75% and the number of silent (synonymous) fixations and near-fixations (greater than 75% frequency). The number of adaptive mutations within each window is calculated as the excess number of replacement (nonsynonymous) fixations or near-fixations above the neutral expectation. Ambiguously sequenced positions (N’s) are ignored. The number of adaptive mutations are normalized by the gene length, and rates of adaptation are calculated as the slope of linear regression fitting adaptive mutations per codon over time. Bootstrap 95% confidence intervals were found by running the same method on 100 bootstrapped datasets. The bootstrapped datasets were created by sampling the codons in the outgroup, with replacement, and then applying the same codon order to the alignment.

Practically, the rate of adaptation is calculated using the `rate of adaptation bhatt.ipynb` notebook inside the `adaptive-evolution-analysis/` directory, which reads in a virus-specific configuration file (in `config/`) that specifies necessary information to complete the analysis as well as metadata about the virus. For instance, the config files specify the relative locations of the necessary input data files, as well as which genes encode the polymerase and receptor-binding protein, whether the virus is enveloped, and what its primary mode of transmission is.

### Rate of adaptation, with an updated outgroup

To account for viruses with especially high rates of evolution where multiple fixations have occurred at the same nucleotide position over the period of time the virus has been sampled, we update the outgroup sequence that is used for computing the rate of adaptation. The starting outgroup sequence is determined as with the ‘fixed outgroup’ method (explained above): as the consensus sequence of all isolates present in the first time window. Then, the outgroup sequence is updated each time a fixation (synonymous or nonsynonymous) occurs. Thus, future time windows are compared to an outgroup sequence that contains information about fixations that occurred in prior time windows. Simply overwriting the outgroup sequence at each fixation event allows more accurate determination of whether future mutations to the same nucleotide site or codon are synonymous or nonsynonymous. However, because this site-counting McDonald-Kreitman based method estimates adaptive mutations in each time window by comparing the alignment to the outgroup, it is essentially counting the accumulation of all mutations that occurred between the outgroup and the time window, with no ‘knowledge’ of whether or not another mutation has previously occurred at any position. This means the method will only ever count a maximum of 1 fixation per nucleotide site. To make the counting method ‘aware’ of fixations that have occurred during previous time windows, the outgroup sequence is stored as a list, with the original outgroup sequence being the first element of the list and fixations getting added as subsequent list elements. At future timepoints, the method is thus ‘aware’ that a fixation has already occurred at any position where the outgroup sequence list has more than one element. The code to implement this method is in the notebook `adaptive-evolution-analysis/rate of adaptation.ipynb`.

### Estimation of threshold, using logistic regression

To estimate the threshold of antigenic evolution, we ran a logistic regression predicting whether or not a virus is evolving antigenically (predictor variable) as a function of the estimated rate of adaptation in the receptor-binding protein (covariate). We used the receptor-binding proteins of three viruses that are known to evolve antigenically (influenza A/H3N2 HA1, influenza B/Vic HA1, and coronavirus 229E S1) and three that are known not to evolve antigenically (measles H, hepatitis A-IA VP1, and influenza C/Yamagata HEF1) in order to have an equal number of viral proteins in both categories for the logistic regression estimation. The threshold rate of antigenic evolution was then obtained as the rate at which the model assigns a greater than 50% probability of antigenic evolution (50% threshold for logistic regression analysis).

### Rate of amino acid substitution

The rate of amino acid substitution was calculated from a phylogeny in order to account for repeated substitutions at the same position. For each virus, we traversed the phylogeny from root to tip, tallying the number of amino acid substitutions that occurred in the receptor-binding protein. Each tip was then plotted according to this accumulated number of substitutions and its sampling date. Linear regression of substitution count and time was used to calculate a rate of amino acid substitution for each virus.

## Supporting information

Supplemental Acknowledgement Table

## Acknowledgements

We gratefully acknowledge the authors who generated and submitted the viral sequences deposited in the NCBI Genbank and ViPR/BV-BRC databases, on which the analyses in this manuscript are based. We also appreciatively acknowledge the authors, originating and submitting laboratories of the SARS-CoV-2 genetic sequences and metadata made available through GISAID Initiative, which supplied the data for the SARS-CoV-2 analyses in this manuscript. We have included an acknowledgements table for NCBI Genbank, ViPR/BV-BRC and GISAID sequences in Supplementary Data. We also thank Allison Li for putting together the norovirus Nextstrain build. Finally, we thank Dr. Harmit Malik for the idea that sparked this project, Dr. John Huddleston for giving it a name, and Cassia Wagner, Dr. Cécile Tran Kiem and Dr. John Huddleston for feedback on the manuscript.

## Supplementary Material

**Figure S1:**
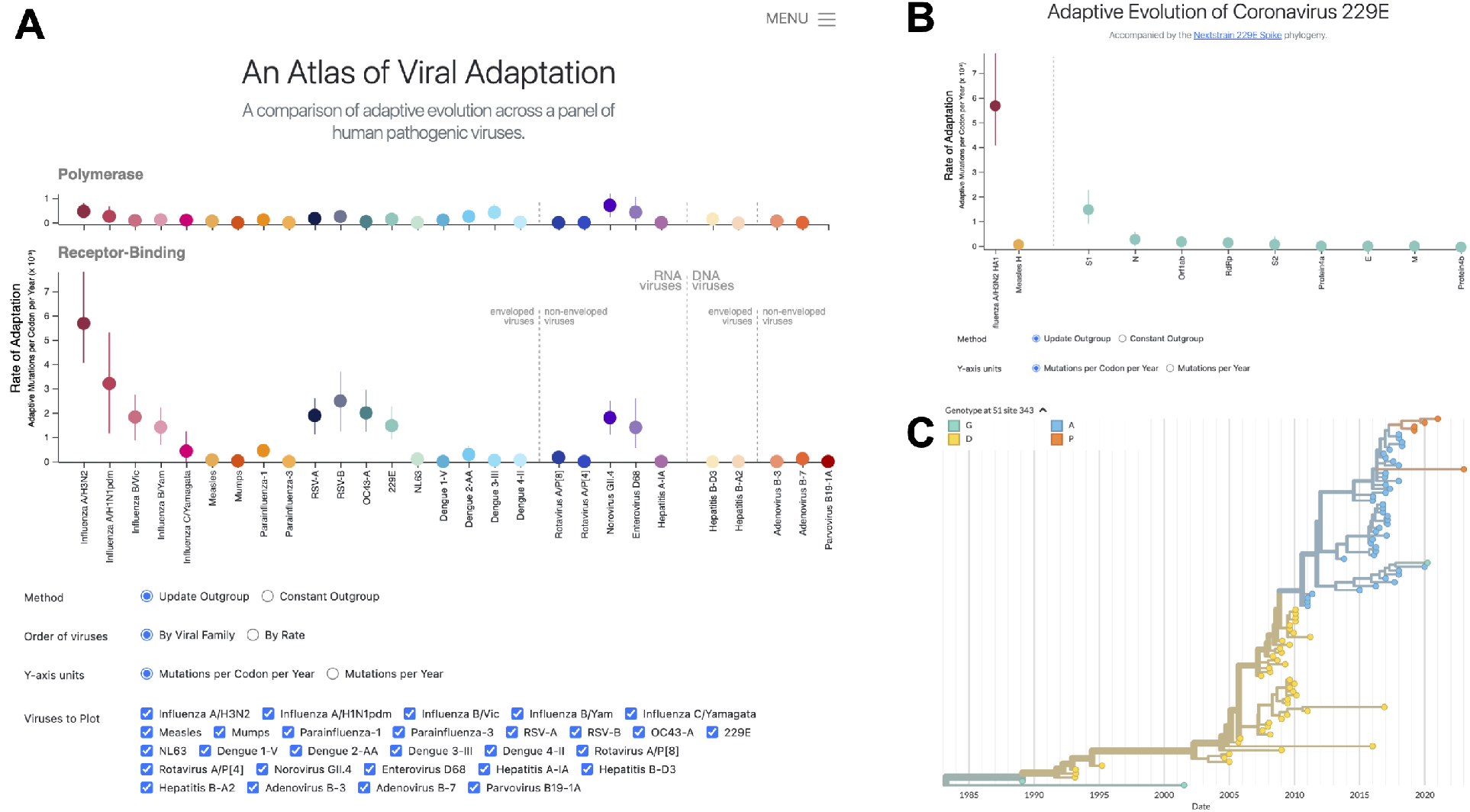
Screenshots of the interactive website presenting the results in this manuscript. **A)** Screenshot of the main page, accessed at https://blab.github.io/atlas-of-viral-adaptation/. Buttons allow the user to toggle the method used to calculate rate of adaptation (update outgroup or constant outgroup), how the viruses are ordered in the plot (by rate or by viral family), the units the rate is displayed in (per year or per codon per year), and which viruses shown on the plot. Hovering over any of the points will display more information about that virus. Clicking on any point will redirect the user to a virus-specific page that shows rates of adaptation across the genome of that virus. **B)** Screenshot of the coronavirus 229E page. **C)** Screenshot of the 229E Nextstrain phylogeny that is paired with the analysis and can accessed from the virus-specific page. In this example, the phylogeny is colored by the genotype at amino acid 343, a residue which has experienced multiple fixation events.

**Figure S2:**
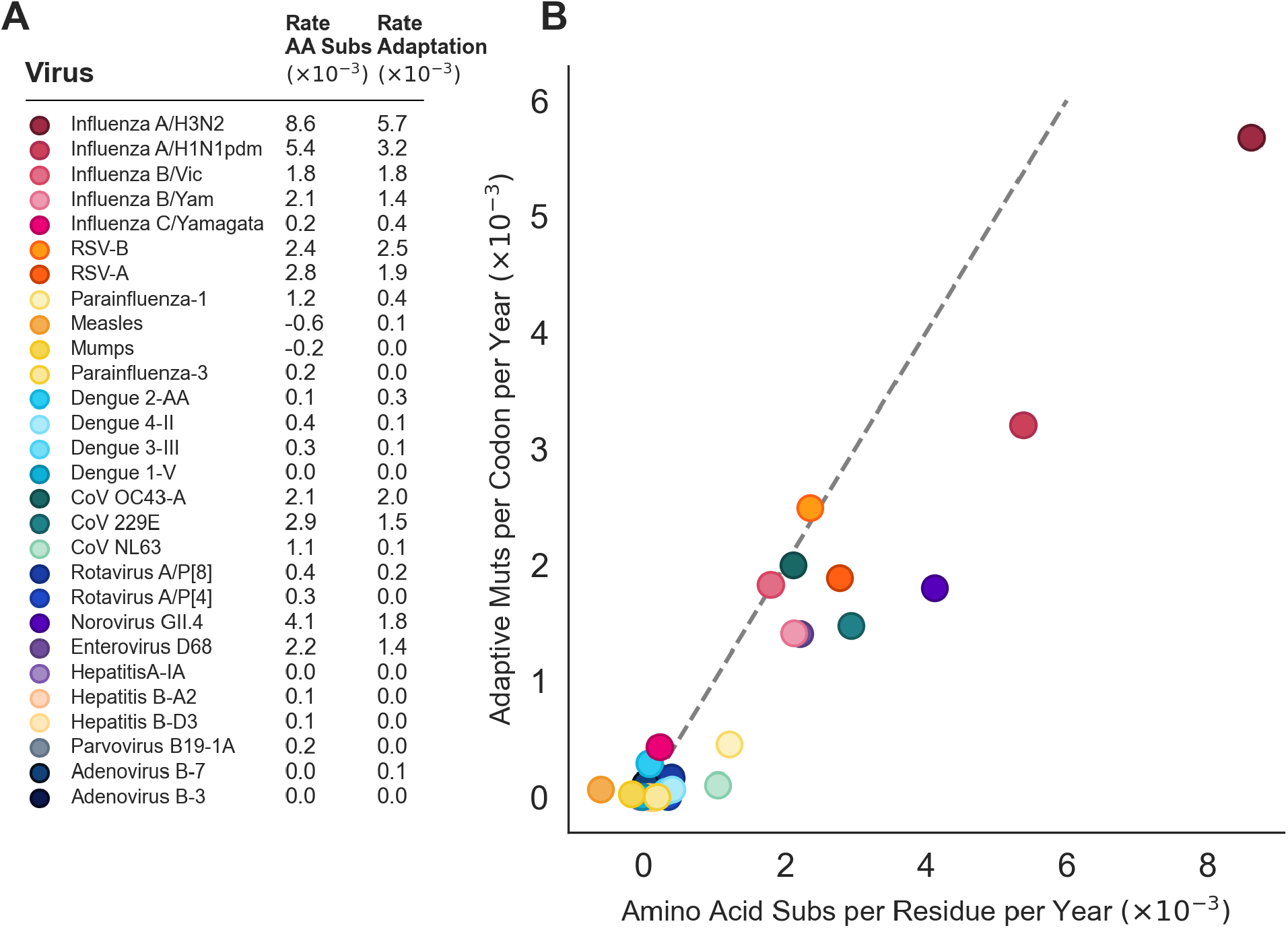
Comparison of rates of amino acid substitution to rates of adaptation. **A)** Rate of amino acid substitution (×10^−3^) and rate of adaptive evolution (×10^−3^) is listed for each of the 28 viruses in the panel. **B)** Rate of amino acid substitution is plotted against rate of adaptive evolution for each virus, with color corresponding to the panel A. The dashed gray line is drawn at *X* = *Y* to indicate the point where all amino acid substitutions are adaptive.

